# Organic electrochemical transistor as an on-site signal amplifier for electrochemical aptamer-based sensing

**DOI:** 10.1101/2022.07.18.500444

**Authors:** Xudong Ji, Xuanyi Lin, Jonathan Rivnay

**Affiliations:** Department of Biomedical Engineering, Northwestern University, Evanston, IL, 60208, USA; Simpson Querrey Institute, Northwestern University, Chicago, IL, 60611, USA; Department of Psychology, The University of Hong Kong, Pokfulam Road, Hong Kong

**Keywords:** organic electrochemical transistor, E-AB sensor, bioelectronics, biosensors

## Abstract

Electrochemical aptamer-based (E-AB) sensors are typically deployed as individual, passive, surface-functionalized electrodes, but they exhibit limited sensitivity especially when the area of the electrode is reduced for miniaturization purposes. We demonstrated that organic electrochemical transistors (OECTs), electrolyte gated tran-istors with volumetric gating, can serve as on-site amplifiers to improve the sensitivity of single electrode-based E-AB sensors. By monolithically integrating an Au working/sensing electrode, on-chip Ag/AgCl reference electrode and Poly(3,4-ethylenedioxythiophene)-poly(styrenesulfonate) (PEDOT:PSS) counter electrode — also serving as the OECT channel, we can simultaneously perform OECT testing and traditional electroanalytical measurement on E-AB sensors including cyclic voltammetry (CV) and square-wave voltammetry (SWV). This device can directly amplify the current from the E-AB sensor via the in-plane current modulation in the counter electrode/transistor channel. The integrated OECT-based E-AB sensor is able to sense transforming growth factor beta 1 (TGF-β_1_) with 3 to 4 orders of magnitude enhancement of sensitivity compared to that in a single electrode-based E-AB sensor (292 µA/dec vs. 85 nA/dec for OECT vs. single electrode SWV). This approach is believed to be universal, which can be applied to a wide range of tethered electrochemical reporter-based sensors to enhance sensitivity, aiding in sensor miniaturization and easing the burden on backend signal processing.

---

Electrochemical aptamer-based (E-AB) sensor have been widely used in the last two decades to sense a large range of targets from ions (1) and small molecules (2, 3) to nucleic acids (4), proteins (5), and whole cells (6). Interest in E-AB sensors is largely due to the merits of aptamers, including their ease of chemical synthesis, strong and tunable binding with specific analytes, wide applicability of different targets, good thermal/environment stability, fast-production and low-cost (7, 8). Electrochemical detection offers sensitive readout of binding-induced conformation changes, often transduced via changes in electron transfer between an electrode and a redox reporter bonded to the aptamer. E-AB sensors based on a single modified electrode, either thin film electrode or bulk wire electrode, have been used as a standard structure in the research community (9). The aptamer is commonly modified with a thiol group on one end to bond with the electrode, and redox reporter on the other end. Signals are usually transduced using established electrochemical interrogation methods like chronoamperometry (CA) (10), cyclic voltammetry (CV) (11), square wave voltammetry (SWV) (12) and electrochemical impedance spectroscopy (EIS) (13). Although the E-AB sensors have been successfully utilized both *in vitro* and *in vivo* (2, 10), the current sensitivity is limited by the surface area of the electrode that determines the amount of aptamers which generate signals after binding with target. Hence, there is a trade-off between high sensitivity and device miniaturization.

One strategy to increase the sensitivity of single electrode-based E-AB sensor is to create electrode with high surface area through electrochemical alloying/dealloying (14), surface wrinkling (15) or electrochemical nanostructuring (16). However, the upper limit for the enhancement of the surface area and the sensitivity can only be improved dozens of times at most (14). Another strategy to enhance sensitivity is through amplification, typically by implementing a transistor (17, 18). Among different types of transistors, organic electrochemical transistors (OECTs) have gained particular attention (19, 20). An OECT is a three terminal device composed of a gate, drain, and source terminal, where a mixed ionic-electronic conducting material forms the channel between source and drain (21). The channel’s conductivity can be altered by the ion injection/extraction controlled through the gate bias (22). Due to its ion-to-electron converting property, high transconductance and biocompatibility, OECTs have proven attractive as biophysical (23–25) and biochemical (26–30) sensors to sense a large variety of targets with high sensitivity and on-site amplification. Unlike frequently used biorecognition elements such as ion-selective membranes, enzymes, and antibodies, aptamers are rarely used in OECTs and very few studies have reported integrating OECTs with E-AB sensors (31–33). In these studies, a typical arrangement is reported where the Au gate is functionalized by the redox-reporter modified aptamer and the sensor output is considered the shift of the transfer curve of OECT. In this scenario, it is difficult for the OECT to capture the modulation of electron transfer kinetics of the redox reporter, which is altered by the structural modulation of the aptamer upon target binding. While operational, many aptamer-based OECT sensors’ sensing mechanisms are likely due to small changes in impedance (most likely capacitive) in the ionic circuit between gate and channel after target binding, which results in the shift of their transfer curve. As such, the typical sensing mechanism in E-AB sensors is not harnessed in previous OECT devices, limiting their generalizability. As a result, a redesigned device concept, architecture and testing scheme is needed to integrate and characterize the OECT-based E-AB sensors. Such a device should fully utilize established sensing mechanisms while taking advantage of the on-site amplification properties of OECTs.

Herein, we developed an OECT-based E-AB sensor by monolithically integrating aptamer-modified Au working/sensing electrode, on-chip Ag/AgCl reference electrode and Poly(3,4-ethylenedioxythiophene)-poly(styrenesulfonate) (PEDOT:PSS) counter electrode. This device retains th features of both OECT and E-AB sensor, guaranteeing the functionality of both. The operation of the E-AB sensor is based on a typical 3-electrode setup and ensures the applicability of established electroanalytical techniques like cyclic voltammetry (CV) and square-wave voltammetry (SWV), retaining the original sensing mechanism. The conductivity changes of the PEDOT:PSS counter electrode caused by the doping/de-doping process from the ionic current during the operation of the E-AB sensor can be monitored with two additional contact leads which provide the output of the OECT device. In this way, direct amplification of the current in the working electrode (gate, E-AB sensor) to the in-plane current modulation in the counter electrode (OECT channel) can be achieved. As a proof of concept, the OECT-based E-AB sensor is used to sense transforming growth factor beta 1 (TGF-β_1_), which is one of the most important biomarkers during wound healing process (34), with 3*∼*4 orders of magnitude enhancement in sensitivity (290 µA/dec for CV-OECT, 292 µA/dec for SWV-OECT) compared to the bare E-AB sensor (24 nA /dec for CV, 85 nA/dec for SWV), with similar detection limit (*∼*1 ng/mL). This approach is believed to be universal since it can be applied to a wide range of tethered redox-reporter-based electrochemical sensors with various electrochemical interrogation methods to enhance sensitivity and improve device form factor and integration.

## Results

The schematic of the OECT-based E-AB sensor and the testing scheme are shown in Fig. 1a and b. The device is composed of three aptamer-modified Au electrodes, one Ag/AgCl electrode and one PEDOT:PSS electrode, which were monolithically integrated using multiple photolithography, vapor phase deposition and etching process (Fig. S1). From the point of view of E-AB sensor, the Au, Ag/AgCl and PEDOT:PSS electrodes are regarded as the working, reference, and counter electrodes, respectively. In this design, established and accepted electrochemical measurements like CV and SWV can be conducted in the E-AB sensor. From the perspective of OECT, the Au and PEDOT:PSS can be considered as the gate and channel respectively, Ag/AgCl serves as a reference point to set the precise voltage drop at gate/electrolyte interface. The interdigitated drain and source electrodes are patterned underneath the PEDOT:PSS channel which defines a large W/L to boost the channel current modulation, while another Au electrode beside the drain and source electrodes serves as the connection lead for the counter electrode, as shown in the enlarged view in Fig. 1a. Fig. S2 shows the microscope image of the OECT-based E-AB sensor where the Au sensing gates and the on-chip Ag/AgCl reference electrode are both 2 mm × 2 mm. The PEDOT:PSS counter electrode is 240 µm × 240 µm with a channel width of 1080 µm and a channel length of 20 µm. During the operation of the OECT-based E-AB sensor, the traditional electrochemical measurement was conducted in the E-AB sensor with the above-mentioned 3-electrode setup, while the change of in-plane conductivity in PEDOT:PSS can be monitored at the same time by the drain and source electrodes and regarded as the output of the OECT device. Since the characterization of the working electrode is performed in a standard 3-electrode setup, the electrochemistry that happens on the working electrode can be retained with respect to the analysis of a single electrode-based E-AB sensor despite the integration with an OECT. Since the current at the working electrode is equal to the current in counter electrode, which induces the ion injection into the counter electrode that dictates PEDOT:PSS conductivity, we can directly relate the output of the OECT to the current in working electrode in the E-AB sensor (also considered as gate current *I*_*G*_ in the OECT perspective).

**Fig. 1.**
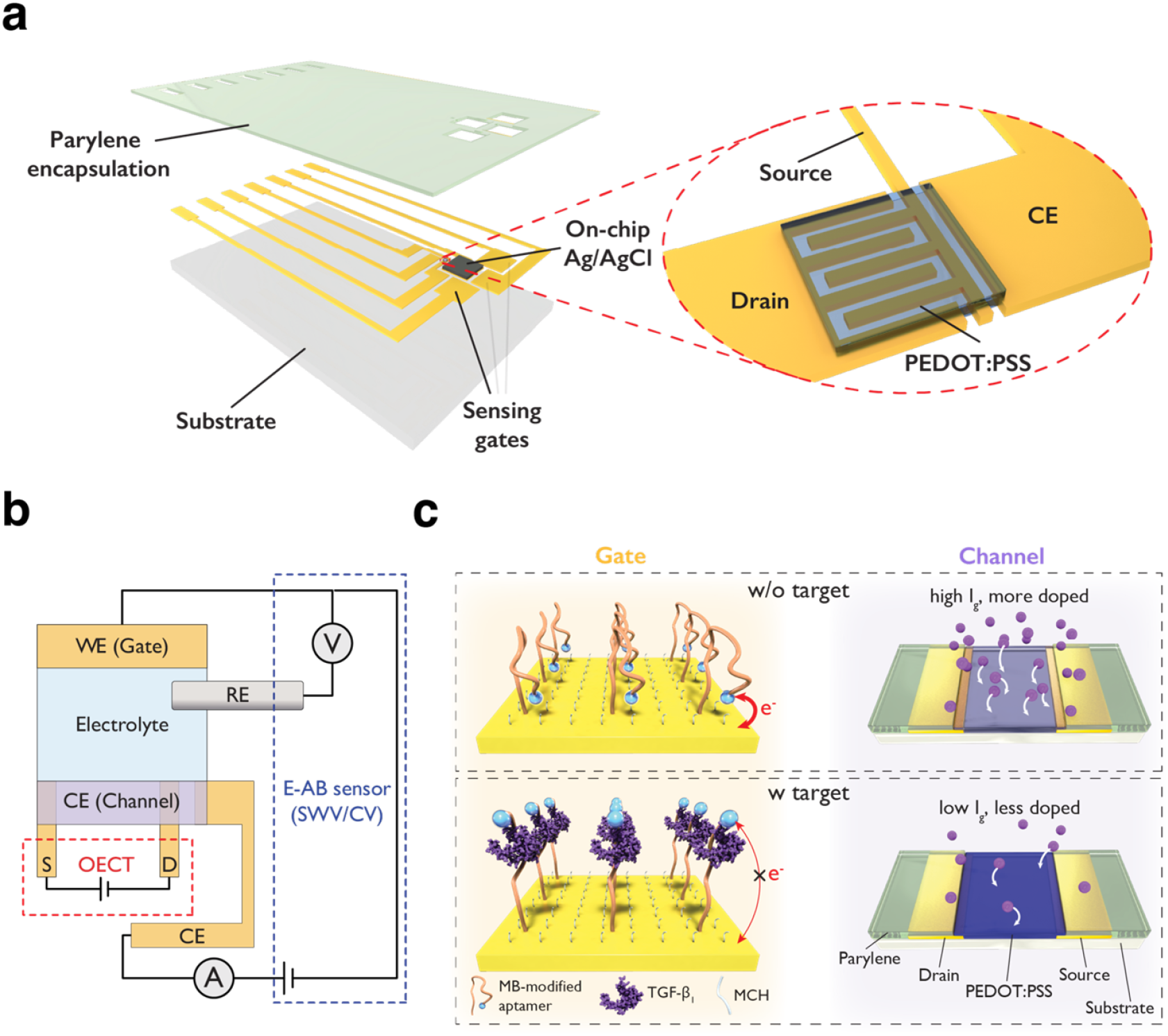
(a) Schematic image of the OECT-based E-AB sensor. (b) Testing scheme of the OECT-based E-AB sensor. Output of channel current in OECT can be monitored during the operation of E-AB sensor in the 3-electrode setup. (c) Sensing mechanism of the OECT-based E-AB sensor for TGF-β_1_. Without the existence of TGF-β_1_, the methylene blue (MB) redox reporter is closer to the gate electrode surface, which results in a large gate current (*I*_*G*_) as well as a larger channel current modulation (*I*_*DS*_). In the presence of TGF-β_1_, a conformational change occurs in the aptamer, and the MB redox reporter moves further from the gate electrode surface, which results in low gate current and smaller channel current modulation.

To demonstrate the proposed sensing mechanism of the OECT-based E-AB sensor, we take the sensing of TGF-β_1_ as an example as shown in Fig. 1c. E-AB sensors for TGF-β_1_ have been demonstrated previously as a “signal-off” type of sensor (35, 36). In brief, in the absence of TGF-β_1_, the aptamer is in a conformation where the methylene blue (MB) redox reporter is close to the Au surface, which results in a large current in working electrode (large *I*_*G*_) during CV or SWV measurement. The large *I*_*G*_ will lead to an elevated ion injection into the PEDOT:PSS counter electrode, which extensively alters the doping level of the PEDOT:PSS, inducing a large channel current modulation recorded by drain/source electrodes. On the other hand, in the presence of TGF-β_1_, the binding between the aptamer and the TGF-β_1_ causes the conformational change of the aptamer and thus moves the redox-active MB away from the electrode surface, which results in a smaller current in the working electrode (small *I*_*G*_). This small *I*_*G*_ will lead to low ion injection into PEDOT:PSS and hence a lower channel current modulation. In this case, the degree of channel current modulation can be regarded as an indicator of the TGF-β_1_ concentration in our OECT-based E-AB sensor.

To support our proposed operation and sensing mechanism of the OECT-based E-AB sensor, we first characterize the aptamer functionalization on the Au electrode, which is a critical step for sensing selectivity and the foundation of the device. The detailed aptamer functionalization process is described in the materials and methods section. X-ray Photoelectron Spectroscopy (XPS) of the aptamer-modified Au electrodes (Fig. S3) shows distinct S 2*p* and N 1*s* peaks originating from the backbone of the aptamer, which indicates the presence of aptamer on the surface of Au electrode. Furthermore, EIS on the Au electrode before and after aptamer modification and mercaptohexanol (MCH) backfill (Fig. S4a) show that the impedance of the Au electrode increased after aptamer modification and MCH backfill over a large frequency range. By fitting the EIS results using Randles circuit, the double layer capacitance on the Au electrode decreased from *∼* 14.1 µF/cm^2^ to *∼* 8.4 µF/cm^2^ after aptamer modification and further to 1.7 µF/cm^2^ after MCH backfill. The EIS results further confirm the successful attachment of aptamer and MCH on the Au electrode. Last, we use EQCM-D to monitor the equivalent functionalization process via mass and electrochemical signal change of the Au-coated QCM sensor (Fig. S5). In Fig. S5b and c, a large mass increase together with a growing redox peak due to MB (attached on the 5’ end of aptamer) in stage 1 indicates the bonding and non-covalent absorption process of aptamer on Au surface. The subsequent mass decrease as well as diminishing of the redox peak in the rising step in stage 2 suggest the removal of the non-bonded aptamer. Next, mass increases again in stage 3, owing to the attachment of MCH, and background signal in SWV results decreases, due to MCH’s ability to insulate the void areas of the Au electrode. Finally, the last rinse step partially removes loosely bound MCH. Combining the results from XPS, EIS and EQCM-D, we can confirm the successful aptamer functionalization on the Au electrode, which is critical for our OECT-based E-AB sensor. By using this established aptamer modification protocol (35), the aptamer-modified Au electrode exhibits stable electrochemical behavior indicated by the consistent SWV results over 30 scans (Fig. S6).

One advantage of our OECT-based E-AB sensor is the monolithic integration of various components in a thin film form-factor, which eliminates the use of bulky reference/counter electrodes and enable future miniaturization and system integration. Functionality of the on-chip Ag/AgCl electrode and PEDOT:PSS counter electrode hence needs to be verified before operating the OECT-based E-AB sensor. EIS of an Au electrode was conducted using either bulky Ag/AgCl pellet or on-chip Ag/AgCl as refence electrode (Fig. S7a), which shows similar results. Then by using on-chip Ag/AgCl as refence electrode, SWV of the aptamer modified Au was also performed with either bulky Pt mesh or thin film PEDOT:PSS as counter electrode, as shown in Fig. S7b. Comparable results were also obtained, which confirm the applicability of PEDOT:PSS as the counter electrode.

With the aptamer-modified Au electrode, functional on-chip Ag/AgCl reference electrode and PEDOT:PSS counter electrode, we demonstrate the operation of the OECT-based E-AB sensor. Simultaneously, we perform CV/SWV operation with working electrode and the OECT measurement as shown in Fig. 2. The detailed voltage waveform and the sampling strategy of the current is schematically shown in Fig. S8. The CV results clearly show the reduction peak of MB in forward scan and the oxidation peak of MB in reverse scan in Fig. 2a. As for the OECT measurement in Fig. 2b, the modulation of the channel current can be observed during the forward and reverse scan of the CV. More importantly, the slope of the channel current change shows a sudden increase when the working electrode undergoes the reduction/oxidation of MB. This is the direct evidence that we can transfer the redox peak information, which is related to the sensing, into the channel current modulation in the OECT. In a similar manner to CV, the SWV and OECT results in Fig. 2c and d also show the conversion of redox peak into the slope change of channel current modulation. Our integrated OECT-based E-AB sensor also exhibits stable operation as indicated by the consistent CV and SWV results among multiple scans as well as the similar channel current changes across the 4 repeated CV and 9 repeated SWV measurements.

**Fig. 2.**
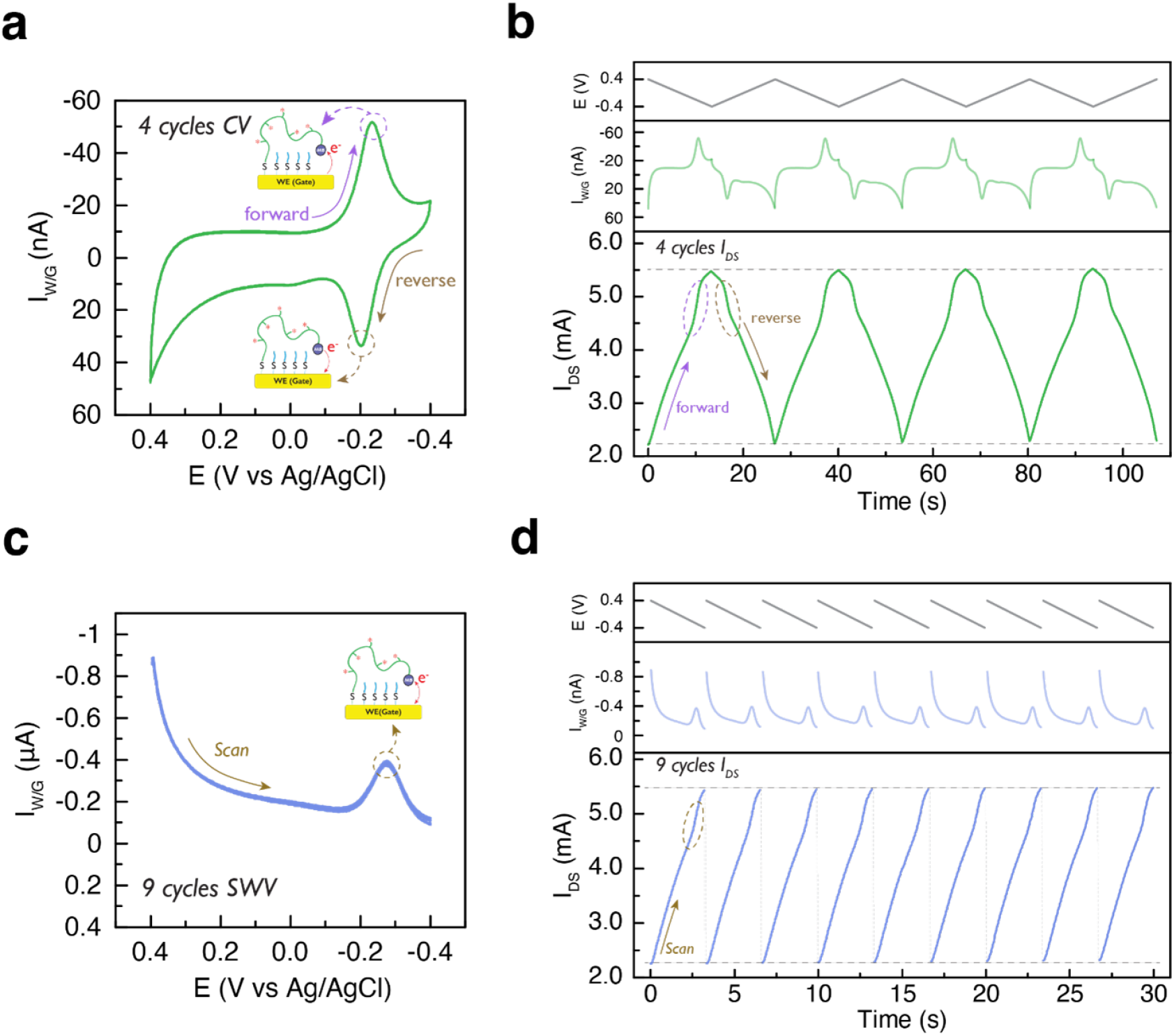
(a) 4 scans of cyclic voltammetry (CV) of the E-AB sensor with scan rate 0.06 V/s and step voltage 0.004 V. (b) Real-time voltage on working electrode, working electrode current/gate current and channel current of the OECT when performing CV measurement in the E-AB sensor. Stable IDS modulation (*V*_*DS*_ =-0.2 V) is demonstrated during 4 CV scans and the sudden change of *I*_*DS*_ indicated by the circles owing to the high reduction/oxidation gate current from the MB redox reporter. (c) 9 scans of square-wave voltammetry (SWV) of the E-AB sensor with scan rate 60 Hz, pulse amplitude 0.04 V and step voltage 0.004 V. (d) Real-time voltage on working electrode (staircase pattern not shown for clarity), working electrode current/gate current and channel current of the OECT when performing SWV measurement in the E-AB sensor. Stable *I*_*DS*_ modulation (*V*_*DS*_ =-0.2 V) is demonstrated during 9 SWV scans and the sudden change of IDS indicated by the circle owing to the high reduction gate current from the MB redox reporter.

Next, we show how our OECT-based E-AB sensor improves the sensitivity compared to the electrode-only E-AB sensor. TGF-β_1_ with different concentrations was added into the electrolyte and the CV, SWV of the E-AB sensor as well as the corresponding OECT channel current modulation were recorded at the same time as shown in Fig. 3. Fig. 3a and d clearly show that the increased concentration of TGF-β_1_ results in a decrease of the redox peak in both CV and SWV, which is coherent with the fact that the binding between aptamer and TGF-β_1_ brings the MB away from the electrode surface and decreases the electron transfer rate. As expected, the decreased redox current in CV and SWV due to the TGF-β_1_ also results in a smaller channel current modulation of the OECT as shown in Fig. 3b and e. A calibration curve can be established when TGF-β_1_ concentration is correlated with the redox current in CV (peak-to-peak current)/SWV (peak current) and the channel current modulation in OECT as shown in Fig. 3c and f. When comparing the sensitivity of E-AB sensor and the integrated OECT-based E-AB sensor for TGF-β_1_ sensing, integrated OECT-based E-AB sensor shows a *∼* 12000-fold enhancement in sensitivity (290 µA/dec) than CV sensing (24 nA/dec) and *∼*3500-fold enhancement in sensitivity (292 µA/dec) than SWV sensing (84 nA/dec) in bare E-AB sensor. In order to demonstrate that the decrease of channel current modulation is indeed caused by the binding between TGF-β_1_ and aptamer rather than the degradation of the PEDOT:PSS channel, stability of the PEDOT:PSS channel was also verified during TGF-β_1_ sensing process (Fig. S9). Standard OECT measurement was performed every time after introducing the TGF-β_1_ with a new concentration by using on-chip Ag/AgCl electrode as the gate. The transfer curves overlapped well with negligible drift, which confirms the stability of PEDOT:PSS channel and further proves that the decrease of channel current modulation in OECT-based E-AB sensor during TGF-β_1_ sensing is indeed caused by the target binding. Finally, this aptamer-analyte binding process was also confirmed by EQCM-D (Fig. S10).

**Fig. 3.**
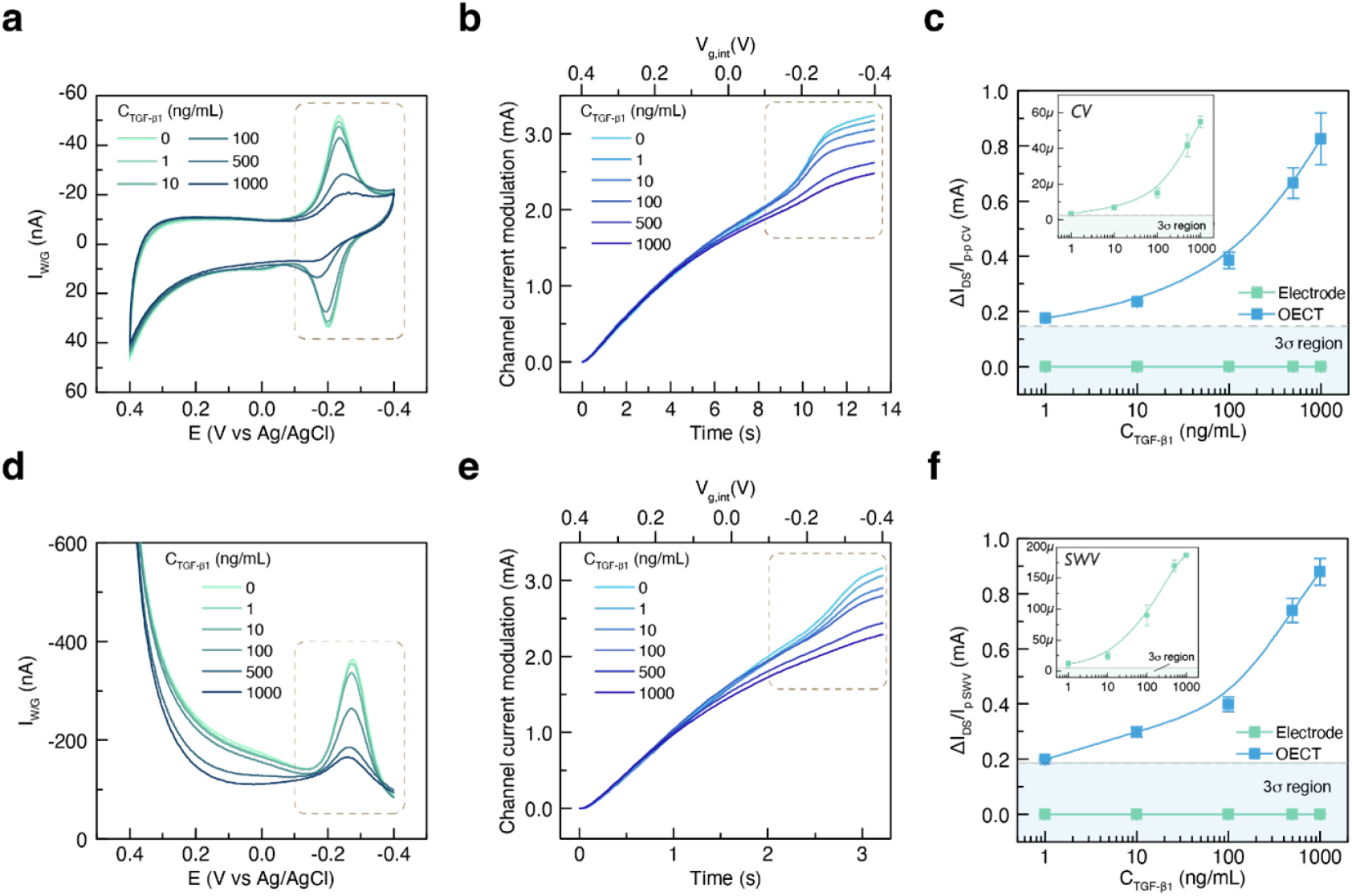
(a, d) Series of CV and SWV results of the E-AB sensor with different concentration of TGF-β_1_. Decreased redox peak is induced by increasing the concentration of TGF-β_1_. (b, e) Series of channel current modulation in OECT with different concentration of TGF-β_1_ when performing the CV/SWV measurement in E-AB sensor. Decreased channel current modulation is induced by increasing the concentration of TGF-β_1_. *V*_*g,int*_ is the voltage at gate/electrolyte interface. (c, f) Calibration curves of TGF-β_1_ sensing based on bare E-AB sensors and OECT-based E-AB sensors (N=3). OECT-based E-AB sensor shows *∼*12000-fold enhancement in sensitivity (290 µA/dec) than CV sensing (24 nA/dec) and *∼*3500-fold enhancement in sensitivity (292 µA/dec) than SWV sensing (84 nA/dec) in bare E-AB sensor. The sensitivity is derived from a linear fitting when the concentration of TGF-β_1_ is larger than 10 ng/mL. 3σ region is defined by the 3-fold of the standard deviation when performing the CV, SWV and the combined CV/SWV-OECT measurement in bare 1XPBS solution for 10 times.

## Discussion

One advantage of our OECT-based E-AB sensor is the decoupling of sensing and amplification. The working electrode for sensing operates in an ideal 3-electrode setup and obeys the original sensing mechanism of a single electrode-based sensor while the mixed-conducting counter electrode/channel is purely used to amplify the current signal from the working electrode. In this situation, our platform is not only useful for transitioning E-AB sensors to OECT-based sensors but is also compatible with other single electrode-based sensors with various electrochemical interrogation techniques, where the current in the working electrode is used as an indicator for sensing.

Typical OECT-based sensors work in a potential-driven mode where the gate voltage is kept as a constant value or scanned in a range and the channel current is monitored, while gate current is often ignored. However, in electrochemical sensing, what really matters is the voltage that has been applied at the gate/electrolyte interface, which drives the electrochemical reaction. However, this voltage at the gate/electrolyte interface is usually unknown in OECT-based sensors. In our device, the introduction of an additional Ag/AgCl electrode serves as an indicator and helps to control the potential drop at the gate/electrolyte interface where the reaction occurs, while the real voltage applied at the channel from the gate is unknown. However, because of the current continuity from working to counter electrode, the gate current is known during the measurement and can modulate the channel current according to the following equation (see Supplementary note 1) (37):

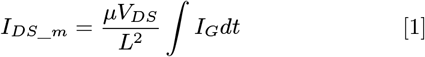

Where *I*_*DS*_*m*_ is the channel current modulation, *L* is the channel length, *µ* is the hole mobility of the mixed conductor and *V*_*DS*_ is the drain/source voltage. The integral of *I*_*G*_ stands for the number of injected ions into the mixed conductor that modulates the carrier concentration during operation. In this scenario, our OECT-based E-AB sensor is a current-driven OECT where the detection of the targets will influence *I*_*G*_ and its integral, hence the channel current modulation. The functionality of equation 1 also helps to explain the observed signals in Fig. 2 and 3. Specifically, single electrode E-AB sensor measured by the CV and SWV show the redox peaks associated with the current in working electrode, equivalent to *I*_*G*_ in the transistor. The transduced signal via the OECT then takes on the functionality of the integral of *I*_*G*_ modified by an amplification factor of 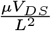. Although the integral of *I*_*G*_ is not used to characterize the single electrode E-AB sensor, it is positively correlated with the redox current in CV (peak-to-peak current)/SWV (peak current). As a result, the amplification factor can still be optimized according to equation 1. It shows that it is highly related to the materials figure-of-merit and the geometry, which indicates that we can further enhance the amplification factor by using organic mixed ionic–electronic conductors (OMIECs) active channel with higher mobility and designing the channel with short length. Although volumetric capacitance (C*), another important parameter of OMIECs, is not explicitly shown in equation 1, it is still critical for the sensor design. When a channel with low volume is used for miniaturization purpose, a larger C* is necessary to maintain the charge injection capacity of the channel, hence its functionality as a counter electrode.

Last but not least, one advantage of OECT-based sensor is the removal of the reference electrode that enables simplicity. While in our device, although we have reference electrode, the thin film form-factor and the monolithically integration enable the minimum added burden in the device design, which still makes this sensor with small size, integration flexibility and easy operation. More importantly, multiple sensing gates that target on different analyte with shared Ag/AgCl reference electrode and PEDOT:PSS counter electrode can be fabricated, which could lead to the multiplexed sensing and amplification on-site.

## Conclusion

In summary, we successfully demonstrated the fabrication and utilization of an OECT-based E-AB sensor that shows 3*∼*4 orders of magnitude enhancement in sensitivity for TGF-β_1_ sensing compared to bare electrode-based E-AB sensor. Monolithic integration of aptamer-modified Au working electrode, on-chip Ag/AgCl reference electrode and PEDOT:PSS counter electrode enables the compact design of the device that has great potential for high density sensing arrays. A new device testing scheme helps to decouple the sensing from E-AB sensor and amplification in OECT, which helps to retain the key features for both OECT and E-AB sensor. This approach guarantees the functionality of both E-AB sensor and OECT and offers an integration strategy between single electrode-based sensor with traditional electrochemical characterization methods and OECT. Direct amplification of the current in the working electrode (gate, E-AB sensor) can be achieved on-site in the OECT, reflected by the channel current modulation. This device concept and testing scheme is believed to be universal for E-AB sensors targeting other analytes, as well as other tethered redox-reporter based sensors, and harnesses well known electrochemical interrogation methods while enhancing sensitivity. In addition, it also allows us to integrate previously discussed high surface area sensing electrodes (14–16), and could further enhance both sensitivity and limit of detection (LOD). As such, these separate techniques are not mutually exclusive, and can be integrated synergistically. We believe this approach can enable compact sensing arrays of aptamer-based sensors that are compatible with various targets, which is critical in the *in vitro* and *in vivo* high throughput diagnostics for trace amount analytes.

## Materials and Methods

### Materials

TGF-β_1_ aptamer was purchased from Integrated DNA Technologies (IDT) with amino modification at the 5 end and thiol modification at the 3 end. The sequence is shown as: 5’/5AmMC6/CG*CTCGG*CTTC*ACG*AG*ATT*CGTGT*CGT TGTGT*C*CTGT*A*C*C*CG*C*CTTG*A*C*C*AGT*C*ACT* CT*AG*AGC*AT*C*CGG*A*CTG/iSpC3//3ThioMC3-D/3’ (35). The backbone of the aptamers was partially modified by phosphorothioate bond (represented by “*” in the sequence) on 5 end of both A and C. This modification is believed to provide enhanced nuclease resistance and higher affinity than the native phosphodiester bond (38). The aptamer is reconstituted at a concentration of 100 µM in IDTE buffer (pH=8.0) from the supplier. Methylene blue (MB), carboxylic acid, succinimidyl was purchased from Biosearch Technologies. PEDOT:PSS (PH-1000) was purchased from Heraeus. All the other chemicals were purchased from Sigma-Aldrich and used as received.

### Aptamer Preparation

NHS-labeled MB was conjugated to the 5’ end of TGF-β_1_ aptamer via the succinimide ester coupling reported previously (36). In short, 50 µL of 100 µM aptamer was mixed with 20 µL dimethylformamide (DMF), 10 µL 0.5 M sodium bicarbonate (NaHCO_3_) and 0.3 mg MB. The mixture was stored at 4 °C for 4 h to modify the aptamer with MB redox reporter. 5 µL MB-modified aptamer was reduced by 10 µL 10 mM tris(2-carboxyethyl)phosphine hydrochloride (TCEP, in IDTE buffer) at room temperature (RT) for 2 h to cleave the disulfer bond in aptamer. This solution was then diluted in 1X Phosphate-buffered saline (PBS) containing 1mM MgCl_2_ to 1 µM aptamer concentration and heated at 95 °C for 5 min to re-fold the aptamer. The aptamer solution was ready to be used for modification after cooling down at RT for 15 minutes.

### Fabrication Process of OECT-based E-AB Sensor

The detailed fabrication process is shown in Fig. S1. First, Cr (5nm)/Au (100 nm) electrodes were patterned on glass substrates by photolithography and the subsequent lift-off process of the negative photoresist (AZ nLOF 2035). After a surface treatment with the adhesion promoter Silane A-174, a 2 µm-thick parylene C was deposited on the substrate serving as an encapsulation layer. Then, a diluted micro-90 2% v/v in DI water) was spin-coated as an anti-adhesive layer, and subsequently, a sacrificial second parylene C layer of 2 µm was deposited. One of the Au electrodes (2 mm×2 mm) was opened through successive photolithography (AZ P4620 photoresist) and reactive ion etching steps (Samco RIE-10NR). 200 nm of Ag was deposited by e-beam evaporator (AJA) and patterned by peeling the sacrificial parylene layer. To form the on-chip Ag/AgCl reference electrode, 0.1 M FeCl_3_ solution was dropped on the patterned Ag electrode for 1 minute to partially transfer Ag to AgCl. After the fabrication of on-chip Ag/AgCl electrode, PEDOT:PSS counter electrode (channel) was patterned using similar sacrificial parylene peel-off process. After opening the exposed counter electrode area, PEDOT:PSS blend consisting of 5 vol% ethylene glycol (EG), 1 vol% (3-glycidyloxypropyl) trimethoxysilane (GOPS), and 0.5 vol% dodecylbenzene sulfonic acid (DBSA), filtered through a 0.45 µm polytetrafluoroethylene filter was spin-coated on the device at 2000 rpm for 1 minute. After a gentle thermal annealing at 90 °C for 2 minutes, the sacrificial parylene layer was removed to pattern the PEDOT:PSS counter electrode (channel), followed by thermal crosslinking at 140 °C for 60 min. The last step of the device fabrication was the modification of the 3 sensing electrodes using TGF-β_1_ aptamer. Same sacrificial parylene deposition and etching process was performed to open the 3 Au working electrodes, while keeping the other part in the device encapsulated to avoid the influence of aptamer on the other components of the device. The device was then immersed in the prepared aptamer solution for 18 h at 4 °C. After rinsing the unbonded aptamer with DI water, the device was then incubated in 2 mM mercaptohexanol (MCH) solution (in 1XPBS with 1 mM MgCl_2_) for 3 h at RT. The device was ready to use after rinsing with DI water and peeling off the sacrificial parylene layer.

### X-ray Photoelectron Spectroscopy (XPS)

The XPS spectrums of aptamer modified Au electrode were taken using Thermo Scientific ESCALAB 250Xi equipped with a monochromatic KR Al X-ray source (spot size around 500 µm) in Northwestern University’s Atomic and Nanoscale Characterization Experimental Center (NUANCE). A flood gun was used for charge compensation. The analysis of the spectrum was performed using the Avantage (Thermo Scientific) software.

### Electrochemical Quartz Crystal Microbalance with Dissipation (EQCM-D)

EQCM-D was performed using an Ivium potentiostat connected with a QSense electrochemistry module. Three-electrode setup comprised an Ag/AgCl reference electrode, Pt counter electrode, and the EQCM chip (Quartz PRO, 5.000 MHz, 14mm Ti/Au) as the working electrode. The mass change was modeled with Sauerbrey equation (39).

### Electrical Characterization

All the electrochemical measurements (EIS, CV, SWV) were conducted using an Ivium potentiostat. OECT channel current was measured by Keithley 2604B source meter with custom-made LabVIEW programs.

## ACKNOWLEDGMENTS

X.J. and J.R. acknowledge financial support from the Defense Advanced Research Projects Agency (DARPA) of the US Dept. of Defense (DOD) through the US Dept. of the Interior under agreement AWD00001593. The content of the information does not necessarily reflect the position or the policy of the Government, and no official endorsement should be inferred. This work made use of the NUFAB and EPIC facility of Northwestern University’s NUANCE Center, which has received support from the ShyNE Resource (NSF ECCS-2025633), the IIN, and Northwestern’s MRSEC program (NSF DMR-1720139).

## Supplementary Information

### Supplementary Note 1

#### Derivation of OECT amplification equation

The channel current can be defined by the Ohm’s Law as:

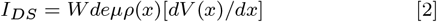

Where *W* and *d* are the channel width and thickness, *e* is elementary charge, *µ* is hole mobility, *ρ*(*x*) is hole density, *V* (*x*) is the voltage along the channel. By assuming the linear increase of the Drain/Source voltage along the channel, the independence of mobility on the hole density, the channel current modulation is purely caused by the change of hole density, which results from the gate current induced ion injection. It can be written as:

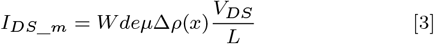

Where *L* is the channel length and ∆*ρ*(*x*) is the change of hole density, which can be defined as:

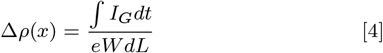

Combining equation 1 ∼ 3, the channel current modulation is given by:

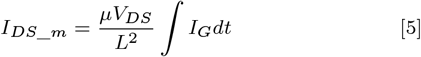

**Fig. S1.**
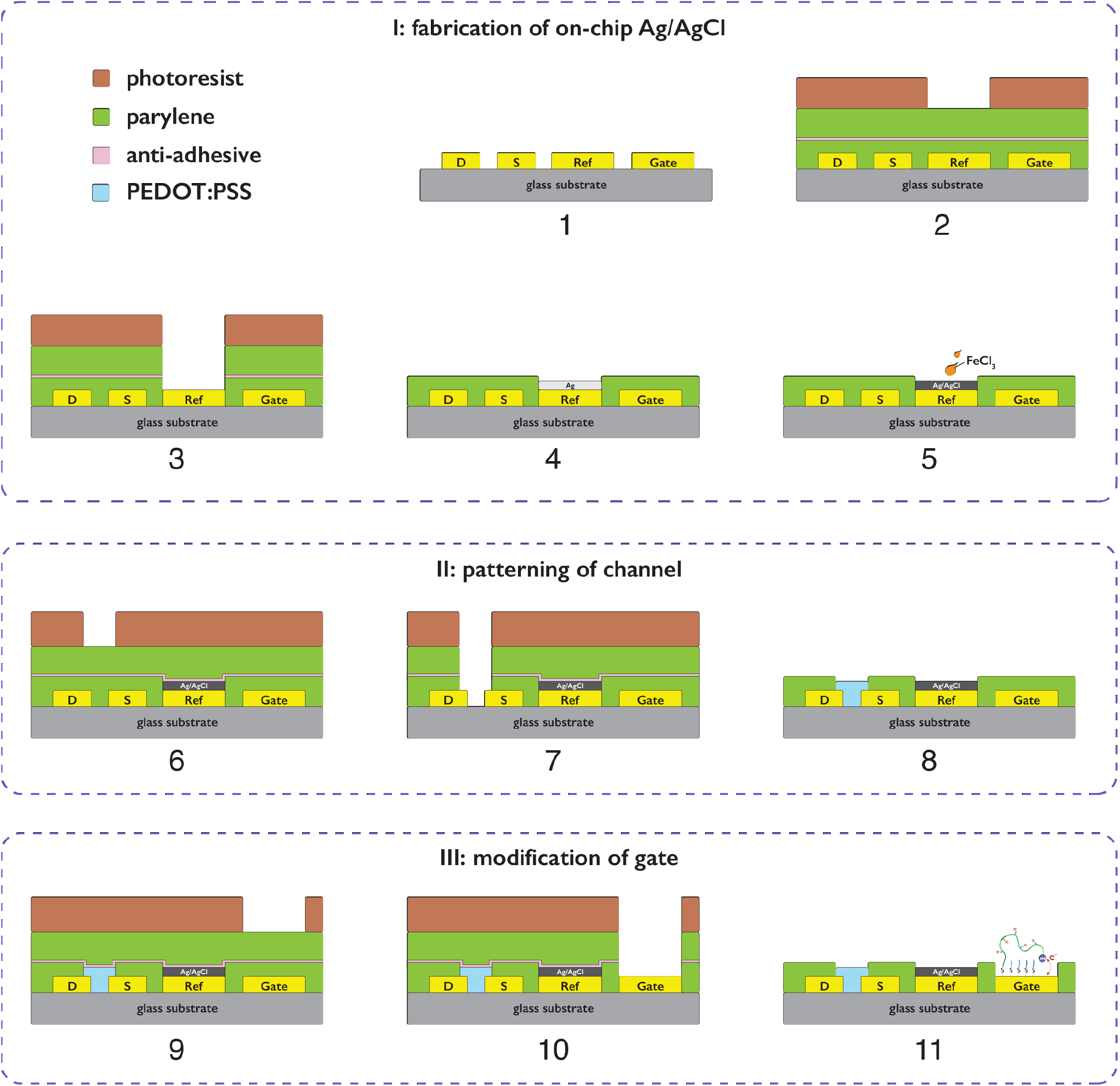
Fabrication scheme of the different components in the OECT-based E-AB sensor. Multiple parylene dry peel-off process have been used to separately pattern the on-chip Ag/AgCl reference electrode, PEDOT:PSS channel and the aptamer modified sensing gate.

**Fig. S2.**
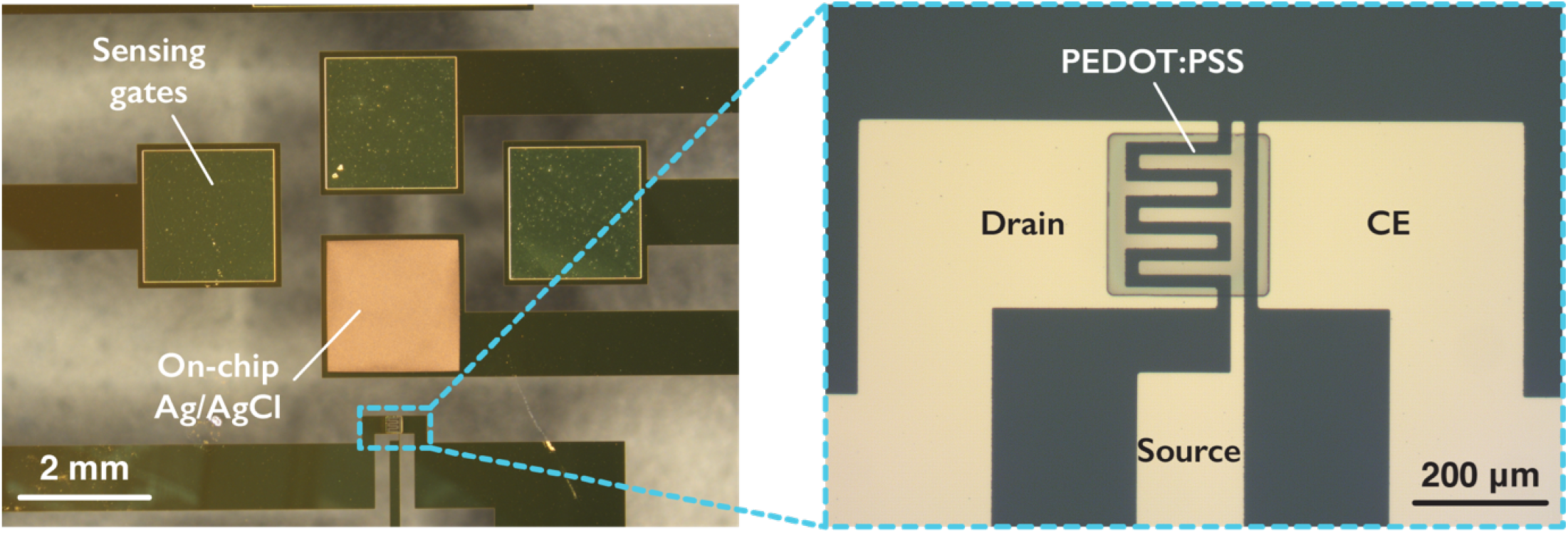
Microscope image of the integrated OECT-based E-AB sensor. 3 working electrodes (sensing gates), an on-chip Ag/AgCl reference electrode and a PEDOT:PSS counter electrode (channel) with interdigitate drain and source electrodes are integrated together.

**Fig. S3.**
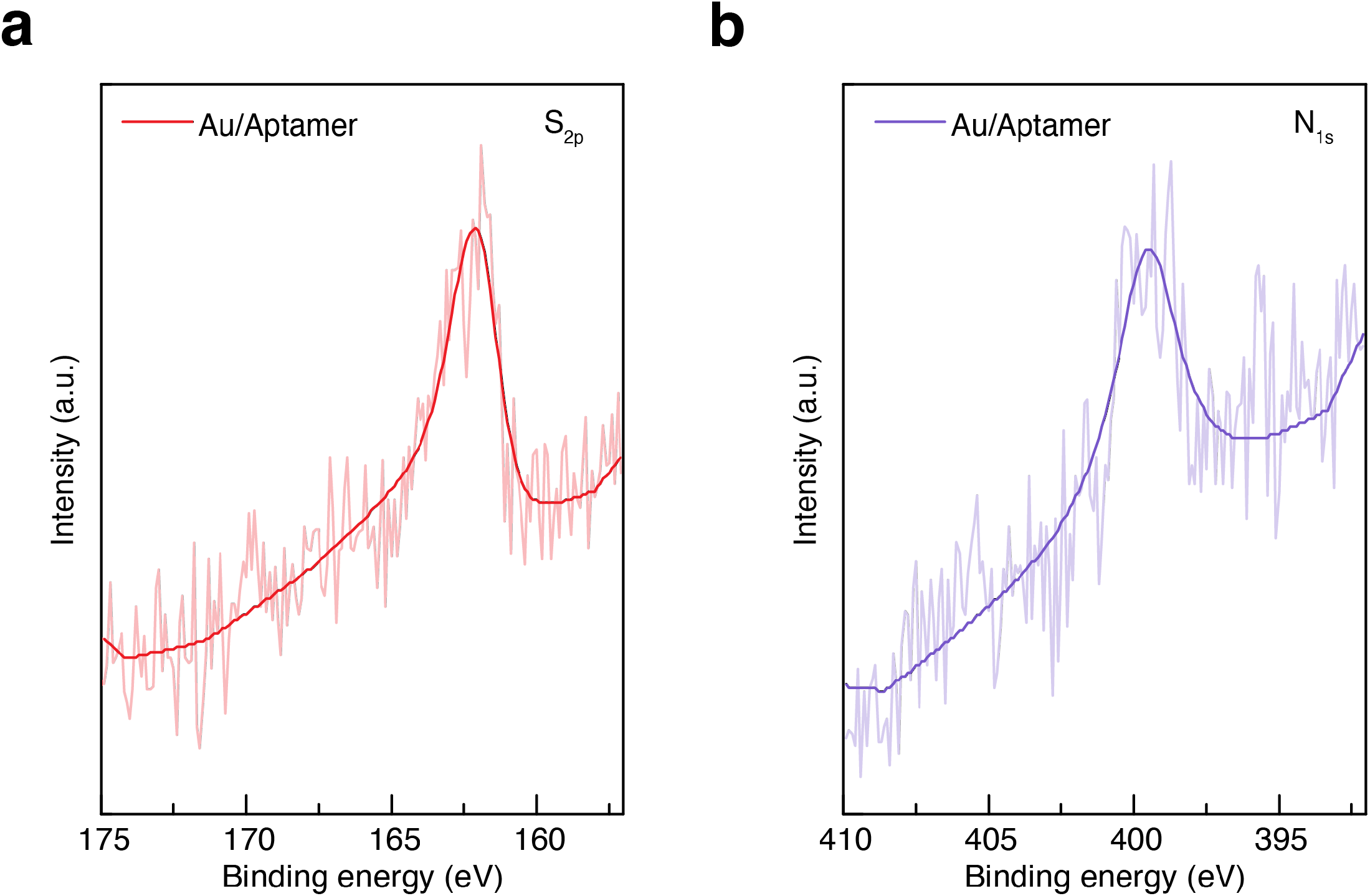
(a, b) XPS of the S 2*p* and N 1*s* of the Au electrode after TGF-β_1_ aptamer modification (after rinse by DI water).

**Fig. S4.**
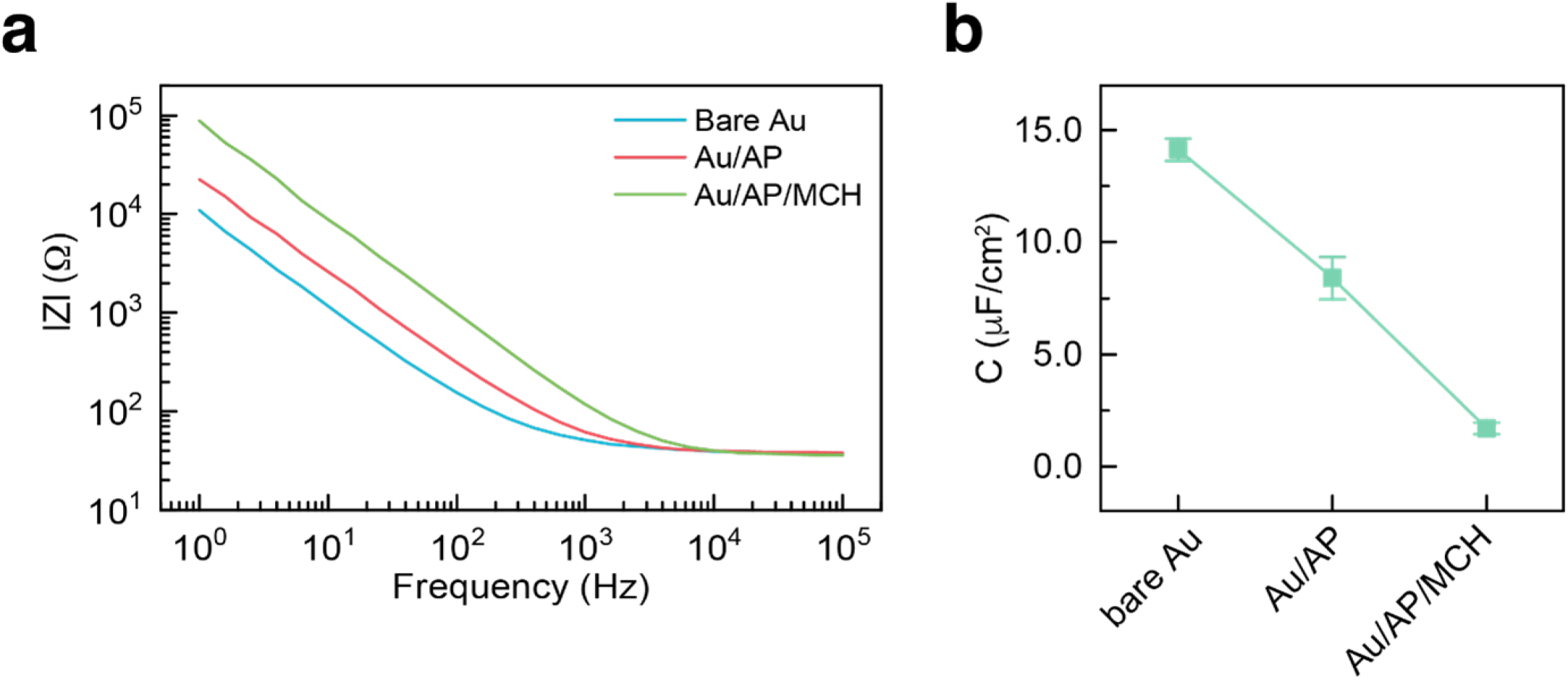
(a) Comparison of the EIS results before and after aptamer/MCH modification. (b) Areal double layer capacitance of the Au electrode before and after aptamer/MCH modification from the fitting of the EIS data.

**Fig. S5.**
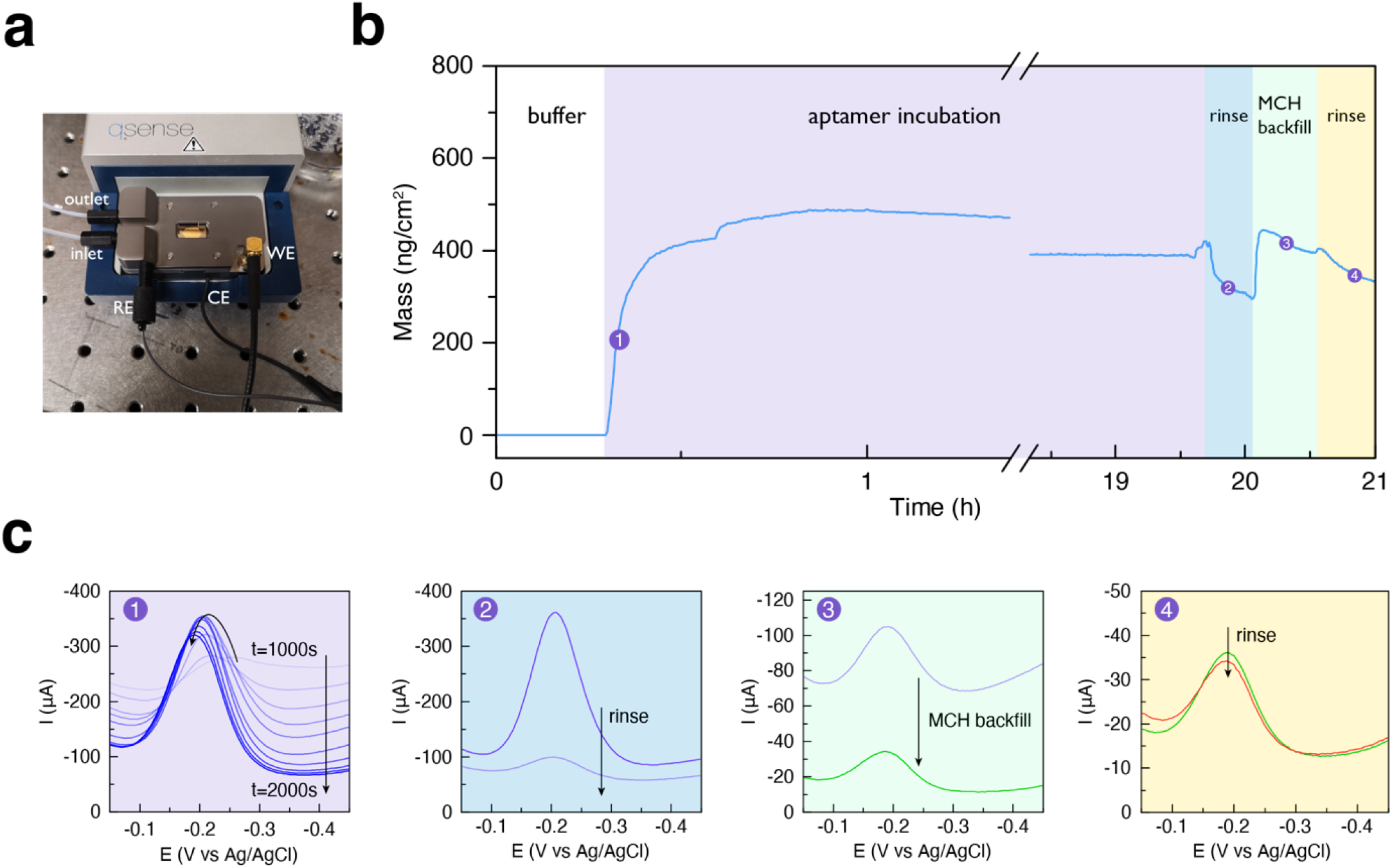
(a) Setup of EQCM-D for characterizing the aptamer modification. (b) Overall mass change on EQCM-D chip during the aptamer modification and MCH backfill. (c) SWV results of the EQCM-D chip during different stages: (1) introduction of aptamer; (2) rinse of aptamer; (3) MCH backfill; (4) rinse of MCH.

**Fig. S6.**
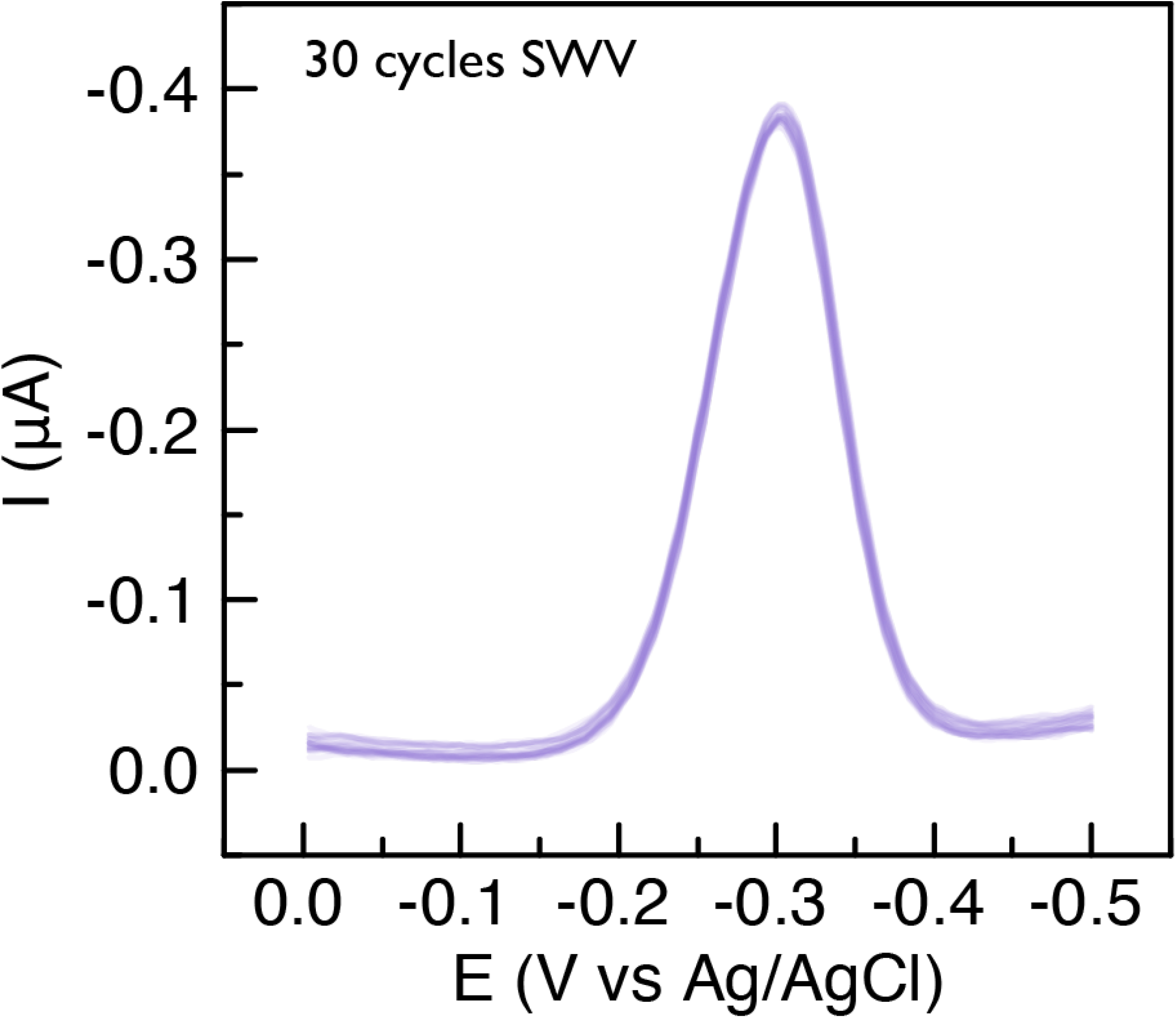
30 cycles of SWV scan overlapped for the aptamer modified Au electrode

**Fig. S7.**
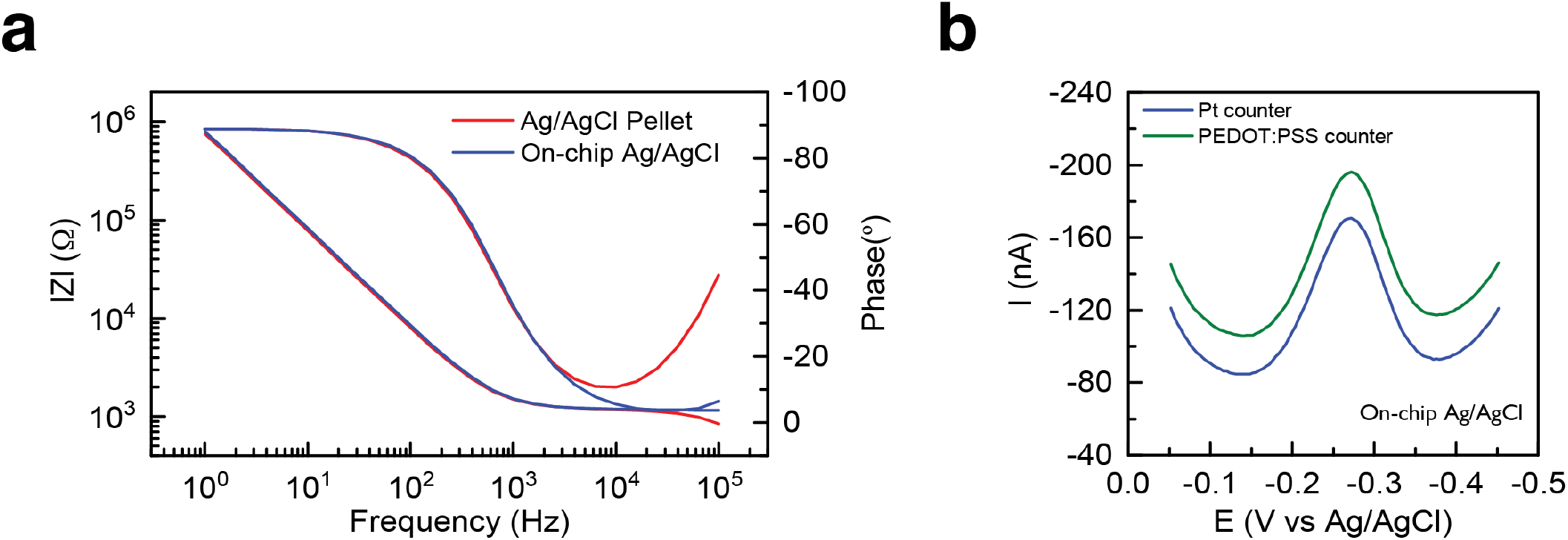
(a) Comparison of the EIS results for an Au electrode with bulk Ag/AgCl pellet and on-chip Ag/AgCl reference electrode. (b) Comparison of the SWV results for an aptamer modified Au electrode with Pt or PEDOT:PSS counter electrode.

**Fig. S8.**
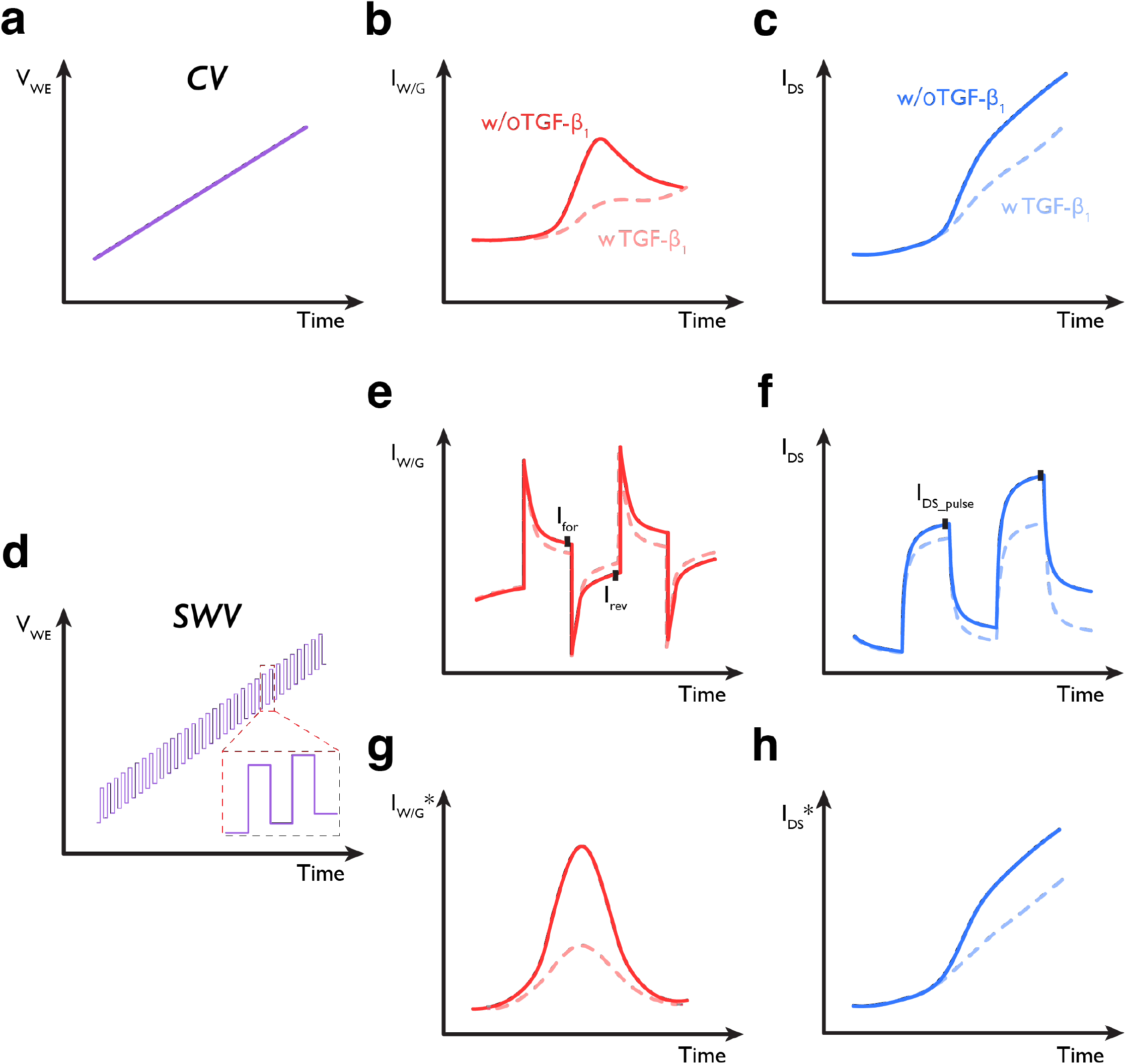
(a) Waveform of CV measurement (only the forward scan has been shown for simplicity). (b, c) Corresponding working electrode current/gate current (defined as *I*_*W/G*_) in *WE* and the resulted *I*_*DS*_ in OECT with or without the existence of TGF-β_1_. (d) Waveform of SWV measurement. (e, f) Corresponding working electrode current/gate current (*I*_*W/G*_) in WE and the resulted *I*_*DS*_ in OECT with or without the existence of TGF-β_1_ (only a short range of voltage pulse similar as enlarged image in Fig. S8d is shown for clarity). The black square is the sampled current that is used to calculate the results in Fig. S8g and h. (g) SWV results in the full voltage range of *WE* with or without the existence of TGF-β_1_. *I*_*W E*_ *∗* = *I*_*for*_ *− I*_*rev*_ for every specific voltage pulse. (h) OECT channel current in the full voltage range with or without the existence of TGF-β_1_. *I*_*DS*_ *∗* = *I*_*DS*___*pulse*_.

**Fig. S9.**
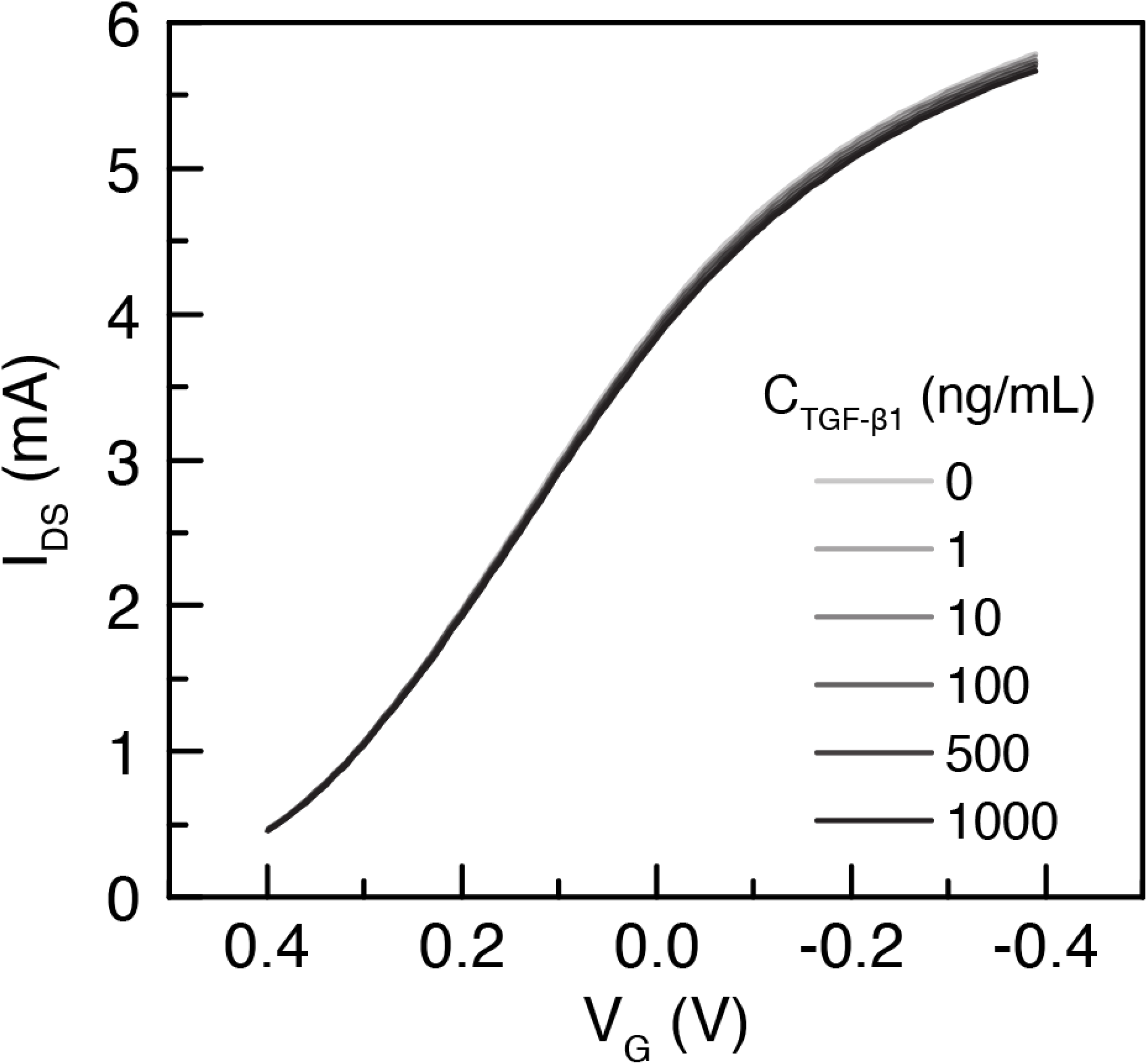
Normal transfer curve measurement of OECT with PEDOT:PSS channel and on-chip Ag/AgCl gate during TGF-β_1_ sensing. Negligible drift of transfer curves can be observed and confirm the stability of PEDOT:PSS channel.

**Fig. S10.**
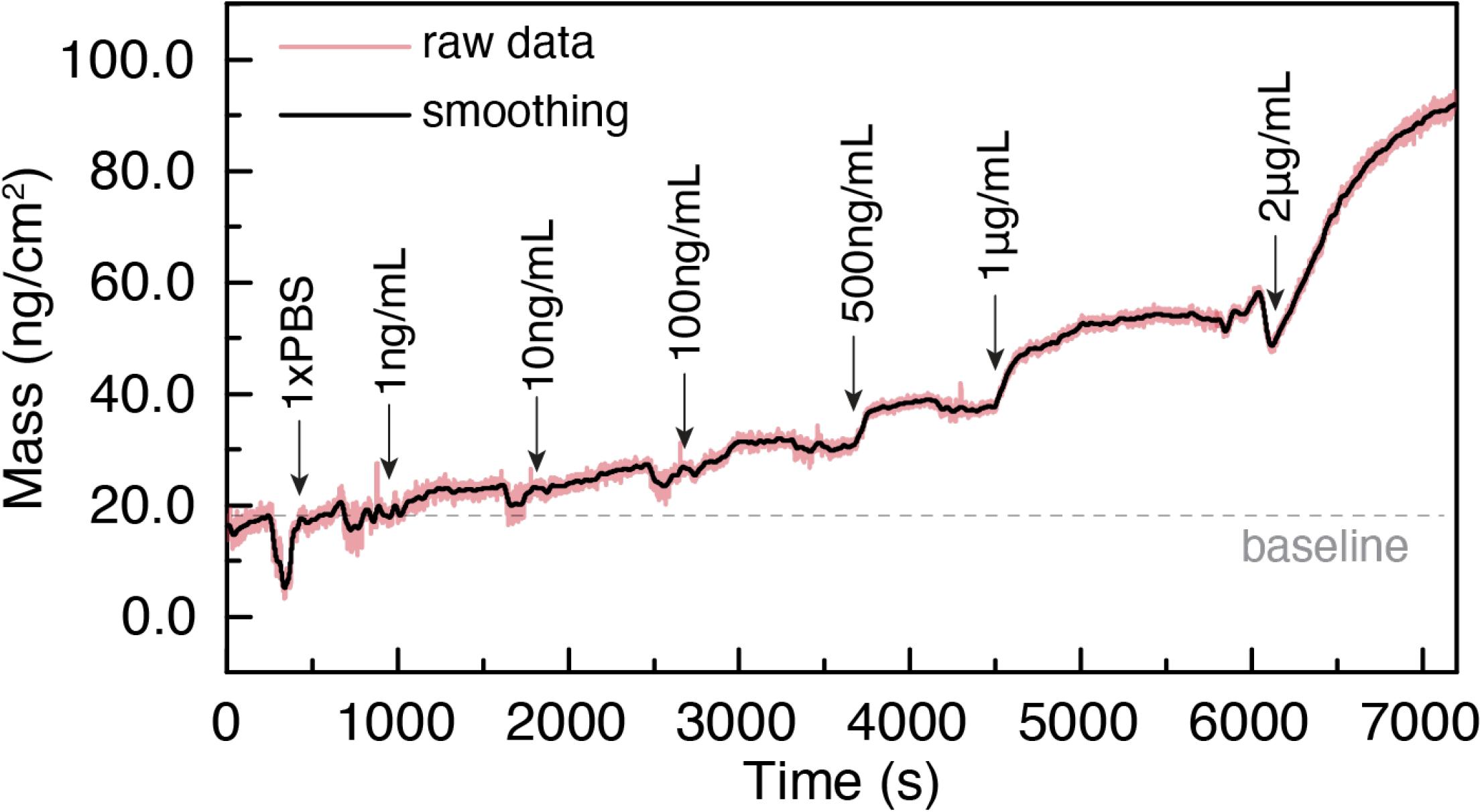
Mass change of the aptamer modified EQCM-D chip after introducing TGF-β_1_ with elevating concentration.

